# From long-term progeny trials to genomic selection: empirical prediction and simulation-guided redesign of Scots pine breeding in Germany

**DOI:** 10.64898/2026.07.21.739892

**Authors:** Bernd Degen, Ana Paula Leite Montalvão, Volker Schneck

## Abstract

Long breeding cycles constrain genetic gain in Scots pine, while mature progeny trials can provide reference populations for genomic selection. We combined phenotypic records from two German trials established in 1990 with dense SNP data. We complemented these data with an offspring-level parentage audit, duplicate filtering, and simulations of breeding strategies. Of 1,986 phenotype-matched genotyped trees, 1,668 formed the parentage-green set and 1,651 remained after exclusion of 17 near-duplicate samples. The final population comprised 1,521 supported controlled-cross offspring and 130 mother-known, father-unknown offspring from 87 progeny labels. Across 14 trait-by-age measurements, GBLUP gave the clearest improvements for diameter and volume from age 20 onwards, whereas height was more mixed. At age 35, ABLUP versus GBLUP heritability was 0.155 versus 0.235 for diameter, 0.265 versus 0.263 for height, and 0.172 versus 0.236 for volume. Leave-progeny-out predictive ability increased from 0.205 to 0.238, 0.249 to 0.265, and 0.203 to 0.239, respectively. SNPscan_breeder simulations compared phenotypic selection, progeny testing, cross-generation genomic selection, and genomic selection with phenotypic thinning under five diversity variants. Progeny testing produced the greatest cumulative gain, but its 45-year cycle reduced annual response. Under the base assumptions, genomic selection with phenotypic thinning gave the highest annual gains for height and fungal resistance, whereas pure genomic selection gave the highest annual diameter gain. Genomic strategies accumulated more kinship than conventional strategies, although a λ = 0.10 kinship penalty improved founder retention with little loss of gain; no mitigation option simultaneously maximised gain, prediction accuracy, and diversity. These results support genomic shortlisting within a field-tested programme with explicit reference-population updating and diversity management.

## Introduction

Scots pine (*Pinus sylvestris* L.) is one of the most widely distributed conifers in Eurasia and remains a major species for timber production and forest restoration across northern and central Europe. In Germany, its silvicultural importance is particularly pronounced on the sandy and comparatively nutrient-poor sites of the north-eastern lowlands, where it has long been a central target of forest tree improvement (Krakau et al. 2013, Schneck 2023). Evidence from established European breeding programmes shows that genetically improved Scots pine reproductive material can provide substantial gains in growth, stem quality and stand yield, and that these biological gains can translate into shorter rotations and improved financial performance (Krakau et al. 2013, Haapanen et al. 2016, Serrano-León et al. 2021). Recent economic analyses likewise indicate that sustained investment in Scots pine improvement can be justified where improved material raises productivity and resilience relative to unimproved planting stock (Saraev et al. 2025).

Conventional Scots pine improvement is founded on plus-tree selection, controlled crossing and long-term progeny testing. Progeny trials provide the basis for estimating additive genetic variation, combining ability and breeding values, but their long duration constrains the rate at which gains can be accumulated and deployed. The Thünen Institute of Forest Genetics and its predecessor institutions established a substantial series of Scots pine progeny tests from the late 1970s onwards. One particularly valuable series was generated from controlled crosses conducted between 1984 and 1987 and planted in 1990 at Eiserbude near Eberswalde, Drewin close to Neustrelitz and Spremberg. After 31 years, the trials still showed marked genetic differentiation in growth and stem form; eleven progenies exceeded the Güstrow seed-orchard control for height by 7.5-13.3%, and height heritability remained sufficiently high to support breeding-value estimation (Schneck 2023). The present study uses the Eiserbude and Drewin/Neustrelitz trials and the current analytical database of 90 progeny labels, comprising 81 controlled-cross progenies, six open-pollinated progenies and three Güstrow seed-orchard reference bulks. These mature trials therefore provide a rare opportunity to connect realised late-age performance with modern genomic information.

Genomic selection offers a means to convert such legacy field resources into reference populations for earlier and more intensive selection. The approach uses genome-wide markers to predict genetic merit without requiring prior identification of individual quantitative trait loci (Meuwissen et al. 2001). For forest trees, its principal attraction is not only the potential to improve prediction, but also the possibility of selecting candidates before mature phenotypes are available and thereby shortening breeding cycles (Grattapaglia and Resende 2011). Genomic relationship matrices replace expected pedigree coefficients with realised relationships, allowing Mendelian segregation among full-sibs to be represented and helping to identify pedigree errors or unrecorded relationships. In addition, single-step genomic BLUP can combine genotyped and non-genotyped individuals within a unified evaluation (Isik et al. 2026). Recent work in large operational pine trials has shown that marker-based relationships can improve genetic parameter estimation and the prediction of individual and family genetic values even when only a subset of trees is genotyped (Tambarussi et al. 2025).

Evidence specific to Scots pine is promising but also demonstrates that the benefit of genomic prediction depends strongly on population design. Calleja-Rodriguez et al. (2020) analysed 694 trees from 183 full-sib families using 8,719 SNPs and found broadly similar prediction efficiencies for genomic and pedigree models, with small advantages for GBLUP and Bayesian ridge regression for several growth and wood-quality traits. Approximately 6,000 markers were sufficient to achieve efficiencies comparable with the full marker set. Importantly, the projected advantage of genomic selection became much larger when a shorter breeding cycle was assumed: halving the cycle approximately doubled relative selection response. However, overall prediction statistics can conceal whether genomic models can rank siblings within families, which is the component of prediction unavailable to pedigree BLUP. In maritime pine, Papin et al. (2024) found similar overall accuracies for ABLUP and GBLUP but near-zero mean within-family genomic accuracy in a training population with too few individuals per family. Their simulations indicated that both total training-population size and the number of genotyped siblings per family are critical. Consequently, random individual cross-validation alone may give an optimistic view of operational performance; leave-family-out, within-family, across-site and across-age validation are required to represent the decisions faced by breeding programmes. Earlier and more frequent phenotyping can complement genomic prediction, as demonstrated by the close agreement between LiDAR-derived and conventional height measurements and selections in a young Scots pine genetic trial (Liziniewicz et al. 2025).

The operational value of genomic selection cannot therefore be judged from one-generation predictive ability alone. Faster cycles and stronger selection can increase gain per unit time, but they may also raise coancestry, increase inbreeding and reduce the representation of founders. Work in Scots pine has shown that useful gain can be retained while maintaining gene diversity when selection is distributed across a sufficiently large number of unrelated families (Fedorkov et al. 2005). Evaluation of breeding strategies must consequently consider annualised response together with inbreeding, kinship, effective population size and founder retention. Stochastic simulation provides a practical way to examine these linked outcomes over recurrent breeding cycles. SNPscan_breeder was developed for this purpose and permits comparisons among phenotypic selection, progeny testing, marker-assisted selection and genomic selection under alternative genetic architectures, mating designs and diversity-management strategies (Degen and Müller 2023a, Degen and Müller 2023b). Calibrating such simulations to empirical heritabilities, prediction performance, family structure and initial relatedness can translate results from the existing trials into decisions about reference-population updating, selection intensity, parent number and mating design.

In this study, we combined long-term phenotypic records from the Eiserbude and Drewin/Neustrelitz progeny trials with dense SNP data and forward simulations of Scots pine breeding. Approximately 2,000 field-tested trees were genotyped with the PiSy50k SNP array (Kastally et al. 2022). The empirical analyses were designed to determine whether these mature trials can serve as a reliable reference population for genomic selection.

Our first objective was to improve the genetic definition of the trial material. We reconstructed latent parental genotypes, evaluated the recorded controlled crosses at the offspring level, and removed near-duplicate samples identified from genome-wide kinship information. This parentage-aware quality control was intended to reduce bias caused by pedigree errors, pollen contamination, sample duplication, or incorrect family assignment.

Our second objective was to compare pedigree-based and genomic prediction methods across all available traits and measurement ages. We compared ABLUP and GBLUP using the same genotyped and parentage-verified trees. Prediction was evaluated with random-individual and leave-progeny-out cross-validation. We also examined site-specific performance, within-family prediction, selection coincidence, and stricter parentage subsets. ssGBLUP was included as a robustness analysis to test whether the joint use of pedigree and genomic relationships improved prediction beyond the separate ABLUP and GBLUP models.

Our third objective was to assess the implications of the empirical results for the future design of the German Scots pine breeding programme. The observed heritabilities, prediction results, relationship structure, and operational time requirements were used to parameterise forward simulations with SNPscan_breeder. These simulations compared phenotypic selection, progeny testing, cross-generation genomic selection, and genomic selection followed by phenotypic thinning. They also evaluated alternative approaches to controlling inbreeding and maintaining genetic diversity.

## Material and method

### Experimental study, plant material and field trials

The study was based on two long-term Scots pine (*Pinus sylvestris* L.) progeny trials of the German breeding programme. Controlled crosses were made between 1984 and 1987, and two-year-old seedlings were planted in 1990 at Drewin near Neustrelitz and Eiserbude near Eberswalde (Schneck 2023). The contemporaneous Spremberg trial was not included because harmonised phenotypic and genomic data were not available for the present analysis.

At each analysed site, the progenies were arranged in three randomised blocks at a spacing of 2.0 × 0.5 m. The current database distinguished 90 progeny labels: 81 progenies from controlled crosses, six open-pollinated or mixed-pollen progenies with a known mother, and three Güstrow seed-orchard reference bulks coded as progenies 88, 89, and 90. Each progeny was represented initially by 60 tree positions per block. The complete experimental grid therefore comprised 90 progenies × 3 blocks × 60 positions × 2 sites = 32,400 tree positions (figure 1).

**Figure 1.**
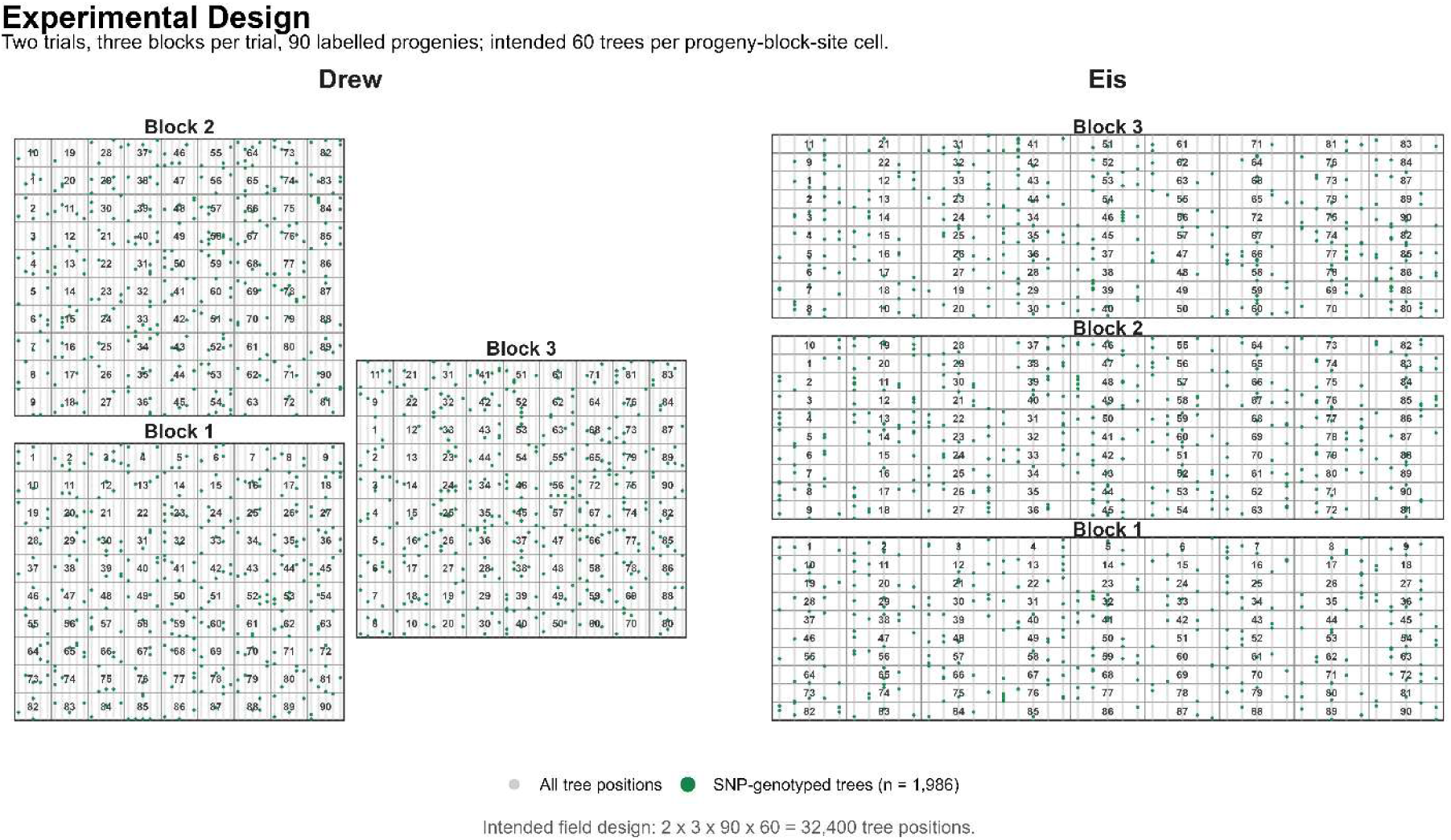
Schematic experimental design of the Drewin and Eiserbude trials. The analysis codes “drew” and “eis” used in the source data correspond to Drewin and Eiserbude, respectively.

Controlled-cross progenies were represented in the pedigree as expected full-sib families. Open-pollinated or mixed-pollen progenies were represented with the recorded mother and an unknown father. The Güstrow reference bulks were treated as background seed lots rather than controlled families. This treatment recognises that open-pollinated or mixed-mating seed lots can contain heterogeneous realised relationships that are not represented adequately by a simple full-sib or uniform half-sib assumption (Araujo et al. 2025, Tambarussi et al. 2025).

#### Phenotypic measurements and data preparation

The full data set comprised 32,400 phenotype rows, 90 progeny records, 58 parent records, and 1,999 DNA-sample mapping records. Repeated measurements covered tree height, stem diameter, individual-tree volume, and stem form (table 1). Measurement ages refer to the nominal ages encoded in the trial database. Stem-form scores were ordinal field assessments but were analysed numerically as exploratory traits; their heritability estimates therefore refer to the observed numeric score scale rather than an underlying liability scale (Gianola 1982).

**Table 1.** Phenotypic variables and numbers of observed records.

| Database code | Trait | Age (years) | Scale | Observed records by age |
| --- | --- | --- | --- | --- |
| h5, h10, h20, h31, h35 | Tree height | 5, 10, 20, 31, 35 | cm | 23,659; 13,204; 11,432; 5,798; 8,042 |
| d10, d20, d31, d35 | Stem diameter | 10, 20, 31, 35 | mm | 13,204; 11,430; 8,033; 8,042 |
| v20, v31, v35 | Individual-tree volume | 20, 31, 35 | m<sup>3</sup> | 11,430; 5,798; 8,042 |
| s20, s31 | Stem-form score | 20, 31 | Ordinal score | 11,405; 8,028 |

*Trait-specific sample sizes declined with age because later inventories included only surviving and measured trees.*

Age-35 survival was calculated relative to the original field positions. Descriptive analyses included site-and progeny-level summaries, growth increments, progeny-mean age–age correlations, and correlations of progeny means between sites. Early-to-late correlations were calculated both after pooling site-specific progeny means and separately within each site to identify confounding by site-level mean differences (Haapanen 2001).

#### DNA-extraction, genotyping and marker quality control

Genomic DNA was isolated from needle tissue according to the protocol described by Bruegmann et al. (2022). The study material was genotyped with the PiSy50k SNP array developed for Scots pine (Kastally et al. 2022). This array comprises 47,712 SNP markers identified from exome-capture data, transcriptomic resources, and resequencing of candidate genes in *P. sylvestris*. Genotyping and preliminary quality assessment were performed at the University of Bristol Genomics Facility, Bristol, UK, and genotype calls were generated using the Axiom Analysis Suite, following the Best Practices Workflow with default settings except sample QC call rate ≥ 93 and average call rate for passing samples ≥ 50. A total of 36014 SNPs were considered “BestandRecommended” and used for downstream filtering and analysis. For genomic prediction, only biallelic SNPs with a marker call rate of at least 0.95 and a minor allele frequency of at least 0.01 were retained. Samples were required to have a call rate of at least 0.90. Following marker and sample filtering, missing genotype dosages were imputed using the corresponding marker mean. The final genomic dataset comprised 1,986 phenotype-matched trees and 34,482 SNPs.

#### KING kinship screen and near-duplicate exclusion

A genome-wide KING analysis was used as an independent genomic quality-control step among the 1,986 matched trees. Pairwise kinship coefficients were estimated with the KING robust relationship inference estimator (Manichaikul et al. 2010), implemented in SNPRelate::snpgdsIBDKING (Zheng et al. 2012), and were used to inspect broad progeny composition and to identify duplicate or near-duplicate genotypes. Screening boundaries were <0.0442 for background or distant relatedness, 0.0442–0.0884 for third-degree-like relationships, 0.0884–0.177 for half-sib or second-degree-like relationships, 0.177–0.450 for full-sib or parent–offspring-like relationships, and ≥0.450 for duplicate or near-duplicate genotypes. These thresholds were used for screening rather than as direct paternity assignments (Manichaikul et al. 2010).

The KING review identified 19 near-duplicate pairs or components and proposed one exclusion per component. Within a recorded progeny, the sample with the higher genotype call rate was retained; for pairs assigned to different progenies, the retained sample was selected from metadata and kinship consistency. Seventeen proposed exclusions occurred in the parentage-green population and were removed before final modelling. Dense marker data therefore provided both pedigree diagnostics and an independent sample-identity control (Tambarussi et al. 2025).

#### Likelihood-based parentage audit

Recorded controlled crosses were evaluated at the offspring level with a 500-SNP likelihood panel selected for high call quality, high polymorphism, and low pairwise LD. Dosages were treated as hard genotype calls when they were within 0.05 of 0, 1, or 2; retained markers had hard-call rate >=0.98, minor allele frequency >=0.10, |r| <= 0.30 after greedy pruning, mean minor allele frequency 0.493, and mean expected heterozygosity 0.500. Mendelian likelihoods used a symmetric genotype-error rate of 0.005.

Because historical parents were not available as genotyped samples, genotypes of 58 named parent labels were reconstructed as latent variables from the 81 recorded controlled crosses using a Mendelian likelihood framework with genotyping error (Marshall et al. 1998, Jones and Ardren 2003, Kalinowski et al. 2007). For each SNP, the joint log likelihood across the connected crossing graph was maximised by iterative conditional updates from 15 starting configurations, with at most 50 iterations per start. The best configuration supplied maximum-a-posteriori parental genotypes, and local conditional likelihoods were converted to posterior-like support probabilities.

Each controlled-cross offspring was then scored against all unordered pairs of reconstructed named parents. The recorded parent pair was accepted for the primary parentage-green set only when it was the highest-likelihood candidate pair. Offspring for which the recorded pair was not top-ranked were assigned to review classes according to whether one recorded parent was retained in the best pair, whether an alternative pair had a top-10 signal, or whether the recorded pair was poorly supported.

Open-pollinated or mixed-pollen progenies were evaluated as mother-known, father-unknown families by comparing named pollen candidates with an unknown-pollen model. The three Guestrow seed-orchard bulk lots were excluded from pedigree-sensitive analyses. The primary parentage-green population retained supported controlled-cross offspring and retained open-pollinated offspring when the unknown-pollen model was preferred. It contained 1,668 genotyped trees before duplicate filtering and 1,651 after removal of 17 KING near-duplicate samples. Two stricter age-35 sensitivity populations were also used: STRICT_IDENTIFIABLE, which retained crosses with sufficiently identifiable parents, and STRICT_EXCLUDE_REVIEW, which additionally removed review crosses (table 2).

**Table 2.** Parentage-aware analytical populations used for the empirical analyses.

| Analysis population | Size | Primary use |
| --- | --- | --- |
| Full phenotypic population | 90 labels; 32,400 positions | Descriptive field-trial summaries |
| Matched genotyped population | 1,986 trees; 34,482 SNPs | Genomic QC and parentage audit |
| Parentage-green before duplicate filter | 1,668 trees; 87 labels | Primary parentage definition |
| PRIMARY_GREEN, final | 1,651 trees; 87 labels | All-trait matched ABLUP–GBLUP analyses |
| STRICT_IDENTIFIABLE | 1,196 trees; 62 labels | Age-35 parent-identifiability sensitivity |
| STRICT_EXCLUDE_REVIEW | 1,137 trees; 58 labels | Age-35 conservative sensitivity |
| Seed-orchard bulk reference set | 61 trees; 3 bulk lots | Excluded from pedigree-sensitive analyses |

*Trait-specific record numbers were lower where measurements were missing. The final PRIMARY\_GREEN population contained 1,521 supported controlled-cross offspring and 130 open-pollinated mother-known offspring from 87 progeny labels.*

#### Pedigree and genomic relationship matrices

For ABLUP, the numerator relationship matrix A was built from the final parentage-audit definition: supported controlled-cross offspring were coded with recorded mother and father, and open-pollinated offspring with recorded mother and unknown father. For GBLUP, the genomic relationship matrix G was calculated from the 34,482 quality-controlled SNP dosages. Matrix summaries and expected-versus-realised relationship comparisons were used as diagnostics before model fitting (Henderson 1976, Van Raden 2008).

For each trait and analytical scope, A and G were aligned to exactly the same parentage-green, duplicate-filtered individuals and phenotype records (table 2). This matched design separates the effect of the relationship matrix from differences in sample composition (Papin et al. 2024, Tambarussi et al. 2025).

#### ABLUP and GBLUP models

The primary relationship-based model was

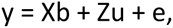

where *y* is the vector of phenotypic observations for one trait at one measurement age, *b* is the vector of fixed-effect coefficients, *u* is the vector of individual-tree additive genetic effects, and *e* is the vector of residual effects. The incidence matrices *X* and *Z* related observations to the fixed effects and individual-tree additive effects, respectively. Additive and residual effects were assumed to be independent (Henderson 1975, Van Raden 2008).

The random terms were specified as

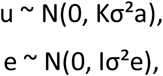

where σ²a is the additive genetic variance, σ²e is the residual variance, and *I* is an identity matrix. The relationship matrix *K* was the pedigree-based numerator relationship matrix *A* for ABLUP or the marker-derived genomic relationship matrix *G* for GBLUP.

Each trait-by-age combination was analysed separately. The repeated measurements at different ages were therefore treated as distinct response variables, and no longitudinal residual covariance or repeated-measures structure was fitted in these models. The analyses comprised five height measurements, four diameter measurements, three individual-tree volume measurements, and two stem-form measurements. Only trees with an observed value for the respective trait and age were included in a given model.

Parentage-aware ABLUP and GBLUP models were fitted with rrBLUP::mixed.solve for the combined two-site data and separately for Drewin and Eiserbude using the rrBLUP r-package (Endelman 2011). The combined analyses contained an intercept and a six-level location-by-block fixed factor representing the two sites and the three blocks within each site. This factor accounted jointly for the mean difference between sites and for environmental variation among blocks within sites. The site-specific analyses contained an intercept and a three-level block fixed factor. Within every trait, age, and analytical scope, ABLUP and GBLUP used identical records and the same fixed-effect design and differed only in the relationship matrix (Endelman 2011, Butler et al. 2023).

The combined and site-specific models can therefore be written as

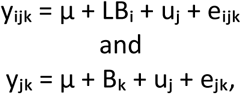

respectively, where μ is the overall intercept, *LB* is the fixed location-by-block effect in the combined analysis, *B* is the fixed block effect in a single-site analysis, and *u* is the random additive genetic effect of an individual tree. No separate random progeny effect was fitted because covariance among trees was represented directly by *A* or *G*. For ABLUP, *A* was constructed from the parentage-audited pedigree: supported controlled-cross offspring were assigned their recorded mother and father, whereas retained open-pollinated offspring were assigned their recorded mother and an unknown father. For GBLUP, *G* was calculated from the quality-controlled SNP dosages.

Variance components were estimated with mixed.solve in the rrBLUP package (Endelman 2011) using restricted maximum likelihood. Individual-tree narrow-sense heritability was calculated as

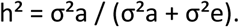

Heritability estimates were reported separately for each trait, measurement age, combined or site-specific scope, relationship model, and analytical population. The estimates represent individual-tree heritability conditional on the fixed site and block effects included in the model. Stem-form scores were ordinal field assessments but were analysed numerically with the same linear mixed-model framework as the continuous growth traits. Their variance components and heritabilities therefore refer to the observed numeric score scale and should be interpreted as exploratory rather than as estimates on an underlying latent liability scale.

#### Site sensitivity

Site sensitivity was evaluated by fitting the same parentage-aware ABLUP and GBLUP models separately to Drewin and Eiserbude and by comparing the resulting heritabilities and cross-validation statistics with those from the combined analysis. In addition, Pearson correlations between progeny means at the two sites were calculated for each trait and measurement age. High correlations indicated comparatively stable progeny performance across environments, whereas low correlations indicated changes in progeny ranking between sites (Burdon 1977, Haapanen 2001).

A full parentage-aware multi-environment variance-partition model was also attempted in ASReml (Butler et al. 2023). The model separated a main additive genetic component shared across sites from a location-specific genetic deviation component. However, the average-information matrix was singular or nearly singular, the common additive component approached the boundary of the parameter space, and nearly all genetic variance was assigned to the location-specific deviation term. The resulting variance-component estimates were therefore considered insufficiently stable for biological interpretation. The final site-sensitivity assessment consequently relied on the separate site-specific ABLUP and GBLUP models and on the observed correlations of progeny means between Drewin and Eiserbude.

#### Cross-validation and secondary prediction analyses

Two matched five-fold validation schemes were used for all traits and scopes. Random-individual cross-validation assigned individuals to permanent folds within location and progeny strata and measured interpolation within the represented population. Leave-progeny-out cross-validation assigned complete progenies to folds so that no member of a validation progeny remained in training data (Papin et al. 2024, Hayatgheibi et al. 2025).

Across the 14 traits, three scopes, two models, and five folds, all 420 random-individual fits and all 420 leave-progeny-out fits completed successfully. Secondary analyses used the cross-validated additive predictions to quantify within-family Mendelian-sampling prediction and ABLUP–GBLUP selection coincidence at 5%, 10%, and 20% selection fractions.

The principal predictive ability was the Pearson correlation between fixed-effect-adjusted observations and cross-validated additive predictions. Observed-scale correlations, RMSE, calibration slopes, fold-level estimates, and validation counts were retained as diagnostics. These empirical phenotype correlations are predictive abilities rather than correlations with unobserved true breeding values (Hayatgheibi et al. 2025).

#### Single-step GBLUP robustness analysis

As a robustness analysis, single-step GBLUP (ssGBLUP) was fitted to the same 1,651 parentage-green, duplicate-filtered trees. The ssGBLUP relationship structure followed the H-matrix formulation that combines pedigree and genomic relationships in a single mixed model (Legarra et al. 2009). The pedigree inverse was generated with ASReml-R, and its genotyped block A_22_ was extracted. G was scaled to the A_22_ scale and blended as G_b_ = 0.90G + 0.10A_22_ to improve compatibility and numerical stability; harmonisation of genomic and pedigree relationship matrices is important for limiting bias in single-step evaluations (Vitezica et al. 2011). The H-inverse updated A^−1^ in the genotyped block by λ(G_b_^−1^ − A_22_^−1^), with λ = 0.5. Positive definiteness of A_22,_ G_b_ and the resulting relationship structure was verified, and individual-tree identifiers and their ordering were checked for exact agreement across the pedigree, genomic, phenotype, and relationship-matrix data before model fitting. The 14 combined-site trait models used the same location-by-block fixed effects as the matched analyses. Observed-scale heritability and the same random-individual and leave-progeny-out five-fold validation schemes were evaluated. Because the ssGBLUP implementation reported correlations of observed phenotypes with genetic predictions, ABLUP and GBLUP predictions were recalculated on the same raw-additive scale for comparison. This analysis tested the integration of pedigree and genomic relationships but did not add phenotype records from non-genotyped trees.

#### SNPscan_breeder simulations

Forward simulations were conducted with the current multi-trait version of SNPscan_breeder, a stochastic, individual-based program developed to compare genome-wide marker-assisted and genomic breeding strategies over recurrent selection cycles (Degen and Müller 2023b, Degen and Müller 2023a). The program represents diploid individuals by SNP genotypes distributed across chromosomes and simulates causal loci, allele effects, recombination, Mendelian segregation, offspring production, phenotypes, selection, mating, and the transmission of genetic diversity across generations.

Trait-specific breeding values were calculated from the effects of the simulated causal loci. Phenotypes were generated by adding environmental deviations to the additive genetic values according to the specified trait heritability. When inbreeding depression was active, the final phenotype was reduced according to

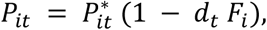

where *P*\**_it_* is the pre-depression phenotype, *F*_i_ is the individual inbreeding coefficient, and *d*_t_ is the trait-specific inbreeding-depression coefficient. In the simulation matrix, *d_t_* = 0.30 for height, diameter and fungal resistance. Thus, an individual with *F_i_* = 0.10 had its phenotype reduced by 3% for each trait. This implementation is a trait-level proportional phenotype penalty; it does not model dominance, recessive deleterious alleles, purging or explicit genetic load.

The three traits were combined in a common multi-trait selection index. Trait values were min– max standardised within the evaluated candidate set as

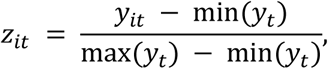

and the index was calculated as

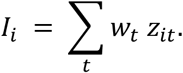

Equal weights were used for the three traits (*w_t_* = 1/3 for height, diameter and fungal resistance). If a raw trait value was outside its specified minimum or maximum threshold, the individual index was set to zero (height: 670–3000; diameter: 43–300; fungal resistance: 0–1000). The upper limits for height and diameter were deliberately widened beyond the current empirical maxima to avoid artificial ceilings during recurrent selection. They therefore functioned as broad plausibility limits rather than as additional economic culling rules. The same index definition was used for all selection strategies within a scenario, ensuring that differences among strategies reflected the information used for selection and the breeding-cycle duration rather than different breeding objectives (Falconer and Mackay 1996, Lynch and Walsh 1998).

#### Population initialization and trait architecture

The simulated population represented an operational Scots pine breeding programme informed by the empirical progeny-test material. Each repetition began with 2,000 candidate individuals generated after a five-generation burn-in. Burn-in used restricted maternal contribution, with 50 mother trees and 2,000 pollen donors. The resulting population had mean kinship approximately 0.013 and tracked ancestor diversity approximately 191, providing broad founder representation with low but non-zero relatedness of the same order as the genotyped trial population (table 3).

**Table 3.** Baseline settings used in the SNPscan_breeder simulations.

| Setting | Baseline value |
| --- | --- |
| Species and application | Scots pine breeding-strategy simulation |
| Candidate population | 2,000 individuals per generation |
| Burn-in | Five generations; 50 mothers and 2,000 pollen donors; mean kinship approximately 0.013; tracked ancestor diversity approximately 191 |
| Simulated traits | height, diameter, and fungal resistance |
| Recurrent selection | Five breeding generations |
| Stochastic repetitions | 30 independent repetitions per scenario |
| Selected reproductive parents | 50 individuals per generation |
| Genome and genomic marker panel | 100,000 SNPs distributed across 12 chromosomes; 10,000 randomly sampled non-causal SNPs used for genomic prediction |
| Cross-generation training proportion | 100% of available reference individuals |
| Phenotypic / progeny-test cycle length | 30 years for PHE; 45 years for PROG |
| Genomic-selection cycle length | 31 years for the first GS or GS+THIN cycle; 16 years for each later cycle |

*The available quantitative-genetic and operational parameters are reported in Supplementary Table S1.*

Three traits were simulated: height, stem diameter, and fungal resistance. The simulated genome comprised 100,000 SNPs distributed across 12 chromosomes. The phenotypic distributions for height and diameter were calibrated using the empirical age-20 measurements as proxies for the traits expressed at the assumed phenotypic selection stage; the corresponding narrow-sense heritabilities were set to 0.50 and 0.25, respectively. Fungal resistance was included as a prospective breeding objective with a heritability of 0.60 and a major-gene component comprising 50 major loci among 1,000 causal loci. This trait was not measured in the two field trials and was therefore interpreted as a hypothetical scenario trait rather than an empirically validated response. All traits used an inbreeding-depression coefficient of 0.30. We did not include pleiotropic effects of causal loci for different traits. Causal effects were generated trait-wise and remained fixed across generations. Complete trait distributions, causal-locus settings, effect distributions, and other configuration parameters are reported in Supplementary Table S1.

#### Genomic prediction and reference-population updating

Genomic selection was implemented with gBLUP using ASReml-R. Predicted genomic breeding values (PGBV) were obtained from the genomic relationship matrix in a mixed-model framework (Henderson 1975, Meuwissen et al. 2001, Van Raden 2008, Butler et al. 2023). The 10,000-SNP marker panel was sampled from the non-causal subset of the 100,000 simulated SNPs, so no simulated causal locus was included directly. Prediction therefore relied on linkage disequilibrium and realised genomic relationships rather than direct observation of causal variants. This provided a less optimistic test, particularly for the hypothetical fungal-resistance trait with major-effect loci. Cross-generation prediction was active for the scenarios GS and GS+THIN: all available reference individuals were used in the baseline training set and the reference information was updated across breeding cycles. The number of candidates, the number of selected reproductive parents, the underlying trait architecture, and all mating and offspring-generation settings were held constant across selection strategies. Thus, comparisons isolated the consequences of the selection information, diversity-management option, and assumed cycle length.

#### Selection strategies

Four selection strategies were compared (table 4). Phenotypic selection and progeny testing represented conventional long-cycle approaches. Pure genomic selection represented direct selection on cross-generation PGBV, whereas genomic selection with phenotypic thinning represented a practical two-stage strategy in which genomic screening reduced the number of candidates requiring subsequent field assessment.

**Table 4.**
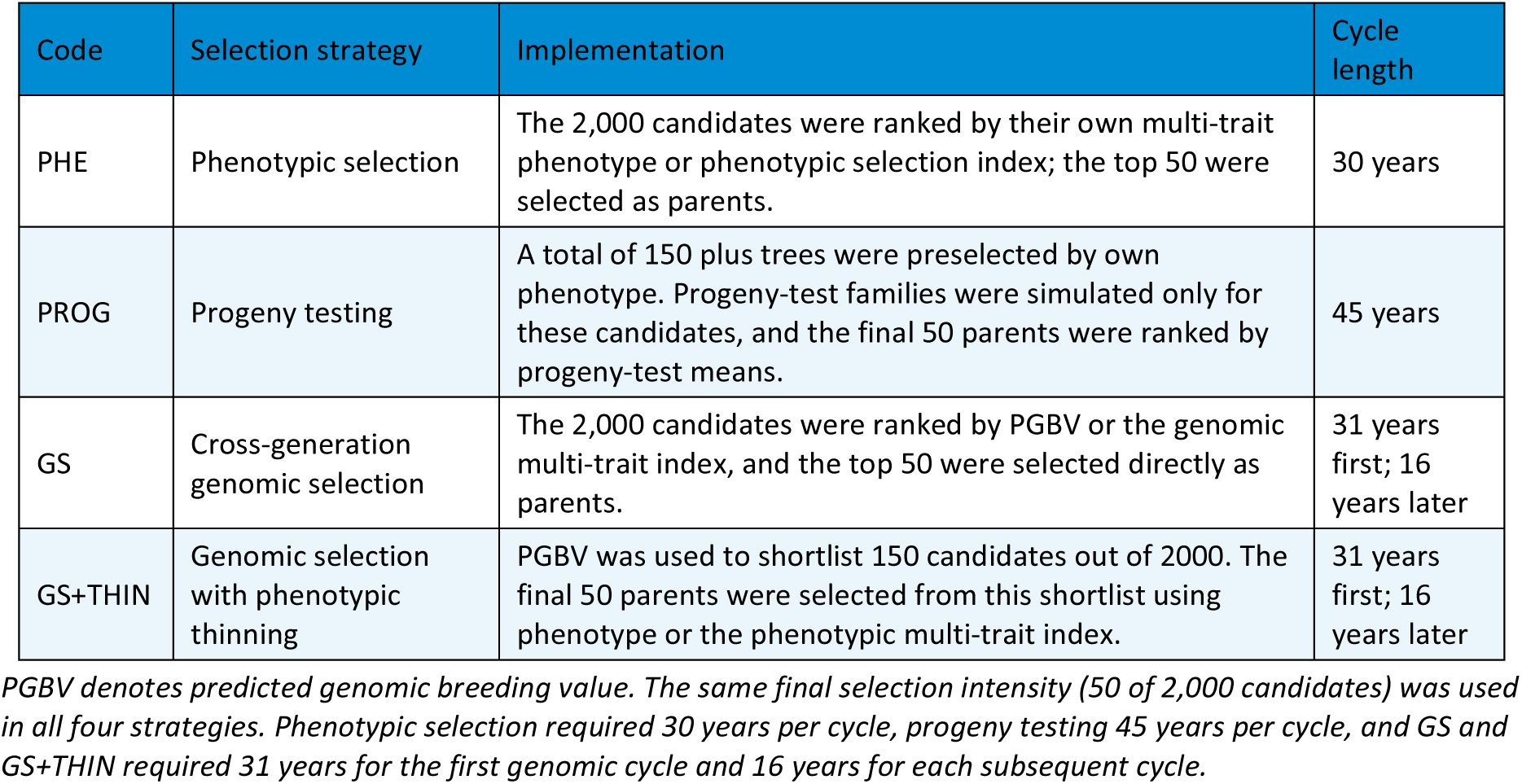
Selection strategies compared in the simulation study.

Phenotypic selection (PHE) ranked all 2,000 candidates by their own phenotypic selection index and selected the best 50 individuals as reproductive parents. Progeny testing (PROG) was implemented as a two-step backward-selection strategy. The 2,000 candidate trees were first ranked by their own phenotypic selection index, and the best 150 were treated as plus trees entering progeny testing (open pollinated progenies – pollen randomly taken from all 2000 candidate trees). Progeny-test offspring were simulated for these plus trees, phenotypes were generated using the same trait architecture and inbreeding-depression settings, and a family mean selection index was calculated. The final 50 parents were selected from the original 150 plus trees according to progeny-test family performance.

Pure genomic selection (GS) fitted ASReml-R gBLUP using the available reference population and ranked the 2,000 genotyped candidates by predicted genomic breeding value or genomic index. The top 50 candidates were selected directly as reproductive parents. Genomic selection with phenotypic thinning (GS+THIN) used the same first genomic-ranking step but retained only the top 150 candidates as a genomic shortlist. Phenotypes were then generated for this shortlist, and the final top 50 parents were selected from the shortlist using the same phenotypic multi-trait index as in the other strategies. The final ranking after thinning used phenotype/index information, not a weighted combination of phenotype and PGBV.

#### Inbreeding, kinship, and gene-flow variants

Each selection strategy was evaluated under five variants, yielding a factorial set of 20 scenario cells (table 5). The base variant included trait-level inbreeding depression without explicit diversity management. The no-inbreeding-depression variant retained the simulated inbreeding and kinship structure but omitted its effect on phenotypes and was treated solely as a sensitivity analysis. The remaining variants tested three mechanisms for limiting relatedness or introducing external genetic material.

**Table 5.** Inbreeding, kinship, and gene-flow variants applied to each selection strategy.

| Code | Variant | Implementation | Interpretation |
| --- | --- | --- | --- |
| BASE | Base model | Inbreeding depression was active; no explicit kinship penalty, unrelated infusion, or external pollen was applied. | Operational reference scenario |
| NO_ID | No inbreeding depression | Inbreeding and kinship were calculated, but trait-level inbreeding depression was not applied to phenotypes. | Biological sensitivity analysis |
| LAMBDA_0.10 | Kinship-penalised selection | The diversity-aware selection option was activated and candidate scores were penalised using a kinship coefficient of $\lambda = 0.10$. | Explicit control of relatedness among selected parents |
| INFUSION | Periodic unrelated infusion | Ten unrelated individuals were introduced from generation 2 at two-generation intervals and replaced the lowest-ranked members of the selected set. Infused individuals entered subsequent genomic reference populations. | Periodic enrichment of the breeding population |
| POLLEN_10 | External pollen | External pollen contributed 10% of mating events in every applicable breeding cycle. | Recurrent gene flow from outside the selected population |

*The principal software settings were UseInbreedingDepression = 1 for BASE and 0 for NO\_ID; GS\_DiversityAwareSelection = 1 and GS\_DiversityKinshipLambda = 0.10 for LAMBDA\_0.10; InfusionStartGen = 2, InfusionInterval = 2, InfusionN = 10, InfusionIncludeInTraining = 1, and InfusionReplaceSelectedAfterRanking = 1 for INFUSION; and UseExternalPollen = 1 with ExternalPollenPercent = 10 for POLLEN\_10.*

#### Simulation execution and annual genetic gain

All 20 combinations of selection strategy and diversity variant were simulated for five recurrent breeding generations with 30 independent repetitions. The same baseline population size, trait architecture, marker number, final number of parents, and number of generations were used throughout. Replicate-specific outputs were retained for every generation so that both the average response and the variability among stochastic repetitions could be compared. Time costs were specified explicitly from the operational sequence represented by each strategy:

a. Phenotypic selection was modelled as forward selection. A total of 2,000 candidates were established from the initial seed source and evaluated at age 15. The best 50 trees were selected to establish a seed orchard, which was assumed to produce seed 15 years later. The resulting cycle length was 30 years.
b. Progeny testing was modelled as backward selection. From 2,000 established candidates, 150 plus trees were preselected from their phenotypes at age 15. Open-pollinated single-tree progenies of these plus trees were then evaluated for 15 years, and the breeding values of the original mother trees were estimated from the mean trait and selection-index performance of their offspring. The best 50 original mother trees were used to establish a seed orchard, which was assumed to produce seed after a further 15 years. The resulting cycle length was 45 years.
c. Cross-generation GS and GS+THIN began by establishing and phenotyping 2,000 trees at age 15 and genotyping them to train the initial genomic-prediction model. Seed collected from the best 50 trees was germinated, and genomic prediction was applied to 2000 of the resulting seedlings at age one. Pure GS selected 50 seedlings directly, whereas GS+THIN selected a genomic shortlist of 150 seedlings for a seedling seed orchard and retained the best 50 on phenotypic or index performance before reproduction. The first genomic cycle therefore required 31 years; once a reference population was available, subsequent cycles required 16 years.

Annual genetic gain was calculated as cumulative percentage gain divided by accumulated strategy-specific years. This is a linear annualisation and not a geometric annual rate.

#### Response variables

Scenario performance was evaluated using response-to-selection, prediction, and diversity metrics (table 6). Trait-specific and index responses were compared both by breeding generation and by elapsed time. For genomic-selection scenarios, prediction accuracy was calculated against the true breeding values generated from the causal loci, which are known in the simulation. Additional phenotype-based correlations were retained to facilitate comparison with empirical cross-validation results.

**Table 6.** Principal response variables obtained from the simulations.

| Response variable | Definition and use |
| --- | --- |
| Cumulative gain | Trait-specific and selection-index response relative to the base population after each breeding generation. |
| Annual genetic gain | Cumulative gain divided by cumulative elapsed years; the primary measure for comparing 30-year phenotypic-selection cycles, 45-year progeny-test cycles, and genomic cycles of 31 years initially and 16 years thereafter. |
| Inbreeding | Mean individual inbreeding coefficient in the selected or reproductive population. |
| Kinship | Mean pairwise kinship among selected or reproductive individuals. |
| Ancestor diversity | Retention and effective representation of tracked founder ancestors across generations. |
| Effective population size | Reproductive and inbreeding effective-size indicators produced by the program. |
| Realised heritability | Generation-specific ratio of realised genetic variance to phenotypic variance. |
| Causal-locus fixation | Numbers or proportions of favourable and unfavourable causal alleles becoming fixed. |
| Genomic-prediction accuracy | Correlation between PGBV and true breeding value, supplemented by correlations with phenotypes for all candidates and for the non-phenotyped or test subset. |
| GS+THIN diagnostics | Mean PGBV and phenotype before and after phenotypic thinning of the genomic shortlist, recorded in GS_Phenotype_Thinning_Records.txt. |

*The principal comparison focused on the joint trajectory of annual gain, inbreeding, kinship, and ancestor diversity* *rather than on genetic gain alone.*

## Results

### Experimental data retention and final analysis population

The cleaned phenotype database contained 32,400 tree positions from two sites, three blocks per site, 90 progeny labels, and 60 initial positions per progeny–block–trial cell. The 90 labels comprised 81 controlled crosses, six known-mother mixed-pollen progenies, and three Güstrow seed-orchard reference bulks. Phenotype availability declined with age: 23,659 height records were available at age 5, whereas diameter, height, and volume at age 35 each had 8,042 records.

A total of 1,986 VCF samples matched phenotype-linked DNA records and passed sample-level filtering, and 34,482 SNPs were retained. The parentage audit identified 1,668 parentage-green records. Seventeen KING near-duplicate exclusions were present in this set, leaving a final primary population of 1,651 genotyped individuals from 87 progeny labels: 1,521 controlled-cross and 130 open-pollinated mother-known records. Detailed counts are provided in Supplementary Table S2.

#### Parentage audit and duplicate filtering

Latent genotypes were reconstructed for 58 parents at all 500 SNPs. Across parent-marker combinations, the mean maximum conditional genotype probability was 0.977; per parent, an average of 430.9 SNPs had support of at least 0.99 and 457.8 had support of at least 0.95, while the lowest parent-level mean maximum probability was 0.917. Among 1,793 controlled-cross offspring, the recorded pair ranked first for 1,536; 61 retained one recorded parent in the best pair, four showed an alternative top-10 signal, and 192 were high-priority review records. The 132 open-pollinated offspring favoured the mother-known, unknown-father model.

The final audit retained controlled-cross offspring only when the recorded pair was top-ranked and retained open-pollinated offspring as mother-known, father-unknown records. The KING duplicate review proposed 19 exclusions overall; 17 occurred in the parentage-green population and were applied before relationship matrices, heritability estimation, cross-validation, and selection comparisons. The final population flow is summarised in Figure 2.

**Figure 2.**
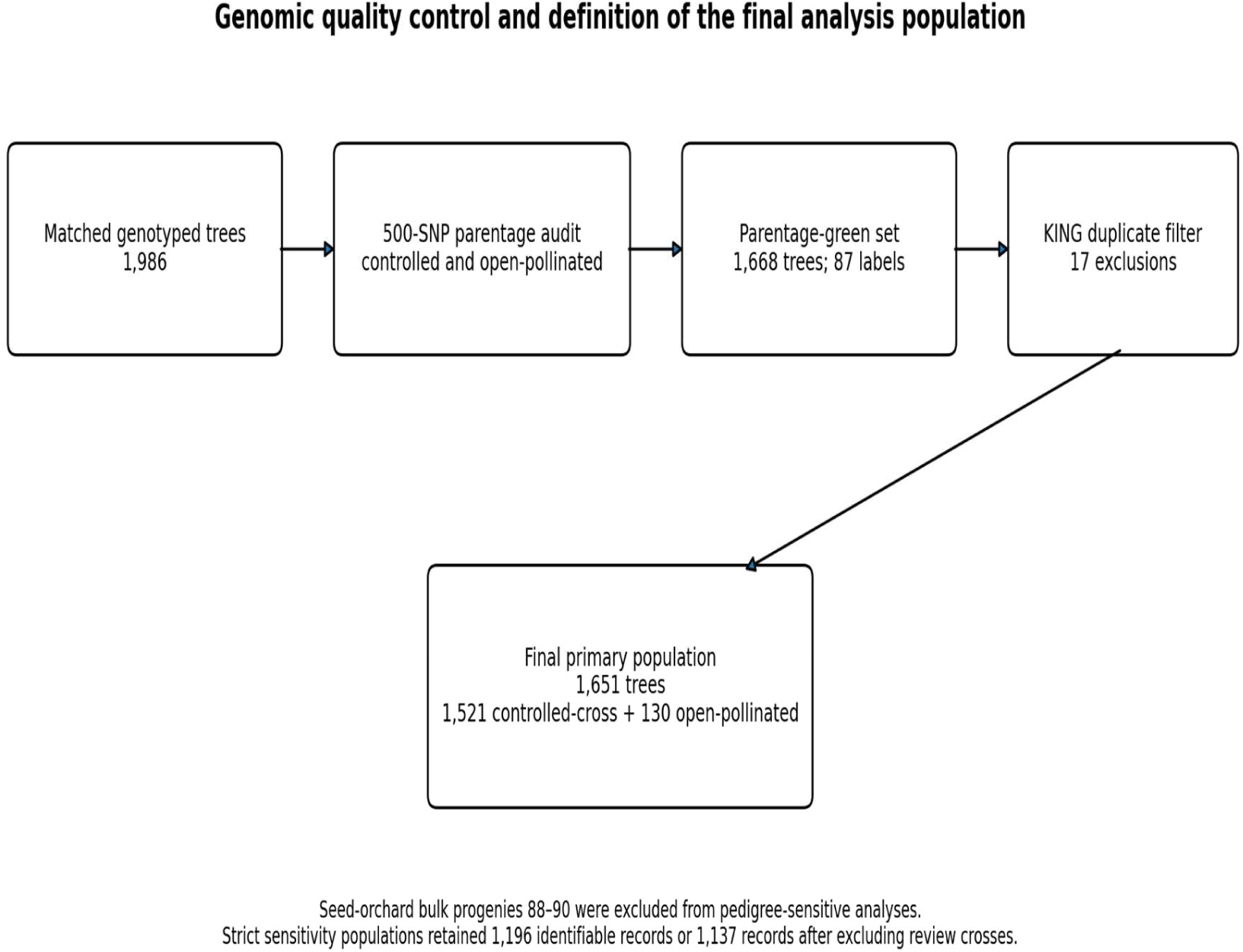
Genomic quality control and definition of the final parentage-green no-duplicate analysis population.

The final relationship matrices were well conditioned for modelling. The A matrix had unit diagonal values, whereas the mean G diagonal was 0.974 and ranged from 0.744 to 1.078; the minimum eigenvalues were 0.500 for A and 0.00086 for G. These diagnostics supported use of both matrices in the matched analyses.

#### Survival, growth dynamics, and youth–age relationships

Age-35 survival was 24.2% in Drewin and 25.5% in Eiserbude. Although the absolute difference was small, screening models detected effects of site, block, and progeny. Diameter, height, and volume increments were generally greater in Drewin, particularly over the age-20 to age-31 interval, confirming persistent site differences in growth.

Progeny-mean relationships strengthened markedly with age. Mid-age diameter and volume were strongly associated with age-35 performance, whereas very early height was strongly confounded by site-level differences. Pooled h5–h35 and h10–h35 correlations were −0.511 and 0.012, respectively, but both were positive within each site. By age 20, the corresponding height correlation was 0.803 across site means, and d20–d35 and v20–v35 correlations were 0.897 and 0.916. The complete set of youth-age correlations is reported in Supplementary Table S3.

#### Longitudinal heritability and predictive ability

Observed-scale heritability was strongly trait and age dependent (Table 7). These estimates were obtained from the combined analysis of Drewin and Eiserbude, with site and block effects accounted for through a fixed six-level site-by-block factor. The model assumed a common additive genetic effect across both sites and did not include a separate site-by-progeny interaction. In the combined primary population, ABLUP was higher for juvenile and mid-age height, with the largest difference at h20 (0.595 versus 0.374). In contrast, GBLUP was higher for diameter and volume from age 20 onwards. At age 35, ABLUP and GBLUP estimates were 0.155 and 0.235 for d35, 0.265 and 0.263 for h35, and 0.172 and 0.236 for v35.

**Table 7.** Combined longitudinal ABLUP and GBLUP heritability and fixed-effect-adjusted predictive ability in the PRIMARY_GREEN no-duplicate population.

| Variable | Age | n | h<sup>2</sup> A / G | Random CV r A / G | Progeny CV r A / G |
| --- | --- | --- | --- | --- | --- |
| h5 | 5 | 1,434 | 0.254 / 0.175 | 0.252 / 0.228 | 0.111 / 0.118 |
| h10 | 10 | 949 | 0.433 / 0.325 | 0.375 / 0.362 | 0.288 / 0.300 |
| h20 | 20 | 1,577 | 0.595 / 0.374 | 0.517 / 0.508 | 0.388 / 0.392 |
| h31 | 31 | 1,173 | 0.469 / 0.374 | 0.474 / 0.482 | 0.400 / 0.414 |
| h35 | 35 | 1,651 | 0.265 / 0.263 | 0.305 / 0.319 | 0.249 / 0.265 |
| d10 | 10 | 949 | 0.162 / 0.133 | 0.186 / 0.164 | 0.144 / 0.147 |
| d20 | 20 | 1,577 | 0.164 / 0.221 | 0.268 / 0.303 | 0.247 / 0.270 |
| d31 | 31 | 1,529 | 0.155 / 0.222 | 0.256 / 0.284 | 0.220 / 0.242 |
| d35 | 35 | 1,651 | 0.155 / 0.235 | 0.248 / 0.286 | 0.205 / 0.238 |
| v20 | 20 | 1,577 | 0.177 / 0.219 | 0.282 / 0.311 | 0.251 / 0.276 |
| v31 | 31 | 1,173 | 0.187 / 0.235 | 0.275 / 0.298 | 0.239 / 0.260 |
| v35 | 35 | 1,651 | 0.172 / 0.236 | 0.257 / 0.290 | 0.203 / 0.239 |
| s20 | 20 | 1,574 | 0.352 / 0.275 | 0.281 / 0.305 | 0.234 / 0.253 |
| s31 | 31 | 1,528 | 0.230 / 0.233 | 0.220 / 0.254 | 0.194 / 0.207 |

Values within each A/G column are ABLUP / GBLUP. Random CV used individual folds; progeny CV withheld complete progenies. Stem-form estimates are exploratory observed-score-scale results.

Random-individual predictive ability showed the same trait-dependent pattern (table 7). GBLUP gains were negative or small for early height and d10, but positive from age 20 onwards for diameter and volume. At age 35, predictive ability increased from 0.248 to 0.286 for d35, from 0.305 to 0.319 for h35, and from 0.257 to 0.290 for v35.

Leave-progeny-out validation was more conservative but retained positive GBLUP deltas for all age-35 target traits: 0.033 for d35, 0.016 for h35, and 0.035 for v35. Positive deltas were also observed for diameter and volume from age 20 onwards and for both stem-form measurements. Complete combined and site-specific heritability and cross-validation results are given in Supplementary Tables S4–S6 and Figures S1–S3.

Stem form did not show a uniform genomic advantage. ABLUP heritability was higher for s20 (0.352 versus 0.275), whereas s31 was similar (0.230 versus 0.233); GBLUP predictive ability was nevertheless higher for both score measurements. The score results refer to the observed numeric scale and should be interpreted as exploratory.

#### Single-step GBLUP robustness analysis

The all-trait ssGBLUP run was numerically stable: A22 and the blended genomic matrix were positive definite, all 14 variance-component models converged, and all 140 cross-validation folds completed. ssGBLUP heritabilities were generally lower than the matched ABLUP and GBLUP estimates; for d35, h35, and v35 they were 0.140, 0.160, and 0.141, respectively. On the common raw-additive prediction scale, ssGBLUP exceeded the better of ABLUP and GBLUP for 6 of 14 traits in random-individual validation and 5 of 14 traits in leave-progeny-out validation, but most differences were small. For age 35, leave-progeny-out predictive ability was 0.246 for ssGBLUP versus 0.245 for GBLUP for d35, 0.246 versus 0.244 for v35, and 0.194 versus 0.225 for h35. In random-individual validation, ssGBLUP was slightly higher for v35 (0.292 versus 0.289), essentially equal for h35 (0.268 versus 0.269), and lower for d35 (0.277 versus 0.289). ssGBLUP therefore did not provide a consistent improvement over the simpler matched models.

#### Strict parentage sensitivity

The age-35 conclusions were robust to stricter parentage definitions (table 8). In the STRICT_IDENTIFIABLE population, GBLUP exceeded ABLUP in heritability and both validation schemes for d35, h35, and v35. The same combined-data pattern remained after additionally excluding review crosses, with the largest gains for diameter and volume.

**Table 8.** Strict parentage sensitivity analysis for age-35 traits in the combined data.

| Population | Trait | n | $h^2$ A / G | Random CV $r$ A / G | Progeny CV $r$ A / G |
| --- | --- | --- | --- | --- | --- |
| Identifiable | d35 | 1,196 | 0.129 / 0.195 | 0.237 / 0.272 | 0.213 / 0.245 |
| Identifiable | h35 | 1,196 | 0.226 / 0.236 | 0.307 / 0.322 | 0.265 / 0.292 |
| Identifiable | v35 | 1,196 | 0.135 / 0.207 | 0.241 / 0.278 | 0.216 / 0.252 |
| Exclude review crosses | d35 | 1,137 | 0.127 / 0.191 | 0.233 / 0.272 | 0.247 / 0.269 |
| Exclude review crosses | h35 | 1,137 | 0.211 / 0.224 | 0.310 / 0.331 | 0.252 / 0.272 |
| Exclude review crosses | v35 | 1,137 | 0.131 / 0.198 | 0.240 / 0.282 | 0.243 / 0.272 |

Values within $h^2$ and CV columns are ABLUP / GBLUP. Both strict populations were filtered for KING near-duplicates before model fitting.

For STRICT_IDENTIFIABLE, GBLUP-minus-ABLUP heritability differences were 0.066 for d35, 0.010 for h35, and 0.072 for v35. After review-cross exclusion, the corresponding differences were 0.064, 0.013, and 0.066. Positive cross-validation deltas in both strict populations show that the main findings were not driven by records with weaker parent identifiability, the corresponding differences are summarised in Supplementary Figure S4.

#### Site sensitivity

Observed progeny-mean correlations between Drewin and Eiserbude were 0.480 for d35, 0.561 for h35, and 0.519 for v35 (table 9). Site-specific estimates showed that the magnitude of ABLUP– GBLUP differences varied between trials, but the clearest genomic advantages remained for diameter and volume. The unstable multi-environment variance partition was not retained.

**Table 9.** Site-specific age-35 heritability and random-individual predictive ability in the PRIMARY_GREEN no-duplicate population.

| Trait | Shared progenies | Cross-site r | ABLUP $h^2$ D / E | GBLUP $h^2$ D / E | ABLUP random CV D / E | GBLUP random CV D / E |
| --- | --- | --- | --- | --- | --- | --- |
| d35 | 86 | 0.480 | 0.179 / 0.130 | 0.270 / 0.173 | 0.245 / 0.197 | 0.278 / 0.213 |
| h35 | 86 | 0.561 | 0.261 / 0.296 | 0.237 / 0.270 | 0.255 / 0.310 | 0.258 / 0.305 |
| v35 | 86 | 0.519 | 0.190 / 0.136 | 0.306 / 0.175 | 0.255 / 0.206 | 0.296 / 0.221 |

Values are Drewin / Eiserbude. Cross-site r is the Pearson correlation of observed progeny means between trials. Complete site-specific random and leave-progeny-out results are provided in Supplementary Tables S4–S6.

#### Within-family prediction and selection coincidence

Within-family Mendelian-sampling prediction was much weaker than the main across-family predictive ability. In combined random-individual validation, within-family correlations were −0.327, −0.353, and −0.322 for ABLUP and −0.011, −0.049, and −0.032 for GBLUP for d35, h35, and v35, respectively. The estimates therefore do not support strong claims about precise ranking of full-sib candidates.

ABLUP and GBLUP additive predictions were nevertheless strongly related. In combined random-individual validation, breeding-value correlations were 0.912–0.918 and top-10% selection overlap was 0.608 for d35, 0.739 for h35, and 0.621 for v35. GBLUP thus produced moderate, trait-dependent re-ranking rather than reproducing the pedigree selection list. Supplementary Figure S5 summarises both analyses.

#### Summary of empirical evidence

The final parentage-aware and duplicate-filtered analyses showed that genomic quality control was a prerequisite for defensible evaluation. GBLUP provided the clearest and most consistent gains for diameter and volume from age 20 onwards, while height remained more similar between models and depended on age and validation design. Strict parentage sensitivity analyses confirmed the mature-trait conclusions. Site-specific results and weak within-family correlations qualify the operational interpretation and support genomic prediction as a complementary ranking tool rather than a replacement for field testing and pedigree evaluation. The ssGBLUP extension was stable but did not consistently improve on the better matched ABLUP or GBLUP model.

### Simulation results

#### Response per breeding generation and per year

Under the BASE assumptions, progeny testing produced the largest cumulative response after five breeding generations for all three traits. Final gains were 17.71% for height, 33.84% for diameter, and 50.51% for fungal resistance. The corresponding gains from phenotypic selection were 11.77%, 23.21%, and 35.00%. Pure genomic selection produced gains of 9.15%, 21.31%, and 28.73%, whereas genomic selection with phenotypic thinning produced 9.33%, 20.49%, and 30.11%, respectively. The two genomic strategies were therefore similar and remained below the conventional strategies in cumulative gain per generation, with small trait-dependent differences between pure genomic selection and phenotypic thinning. Complete scenario values and the corresponding cumulative-gain plot are provided in Supplementary Table S7 and Figure S6.

At generation 5, PHE had accumulated 150 years, PROG 225 years, and GS and GS+THIN 95 years. Under BASE, final annual gains for height, diameter, and fungal resistance were 0.078%, 0.155%, and 0.233% yr−1 for PHE; 0.079%, 0.150%, and 0.224% yr−1 for PROG; 0.096%, 0.224%, and 0.302% yr−1 for GS; and 0.098%, 0.216%, and 0.317% yr−1 for GS+THIN. As visible in figure 3, GS+THIN was highest for height and fungal resistance, whereas pure GS was highest for diameter. Across all 20 scenario cells, the largest annual gains occurred for pure GS under NO_ID: 0.111% yr−1 for height, 0.251% yr−1 for diameter, and 0.332% yr−1 for fungal resistance. Because NO_ID omits the phenotypic cost of inbreeding while retaining the associated relatedness, these values represent an optimistic upper bound rather than an operational recommendation. Annual-gain values for all scenario combinations are provided in Supplementary Table S8.

**Figure 3.**
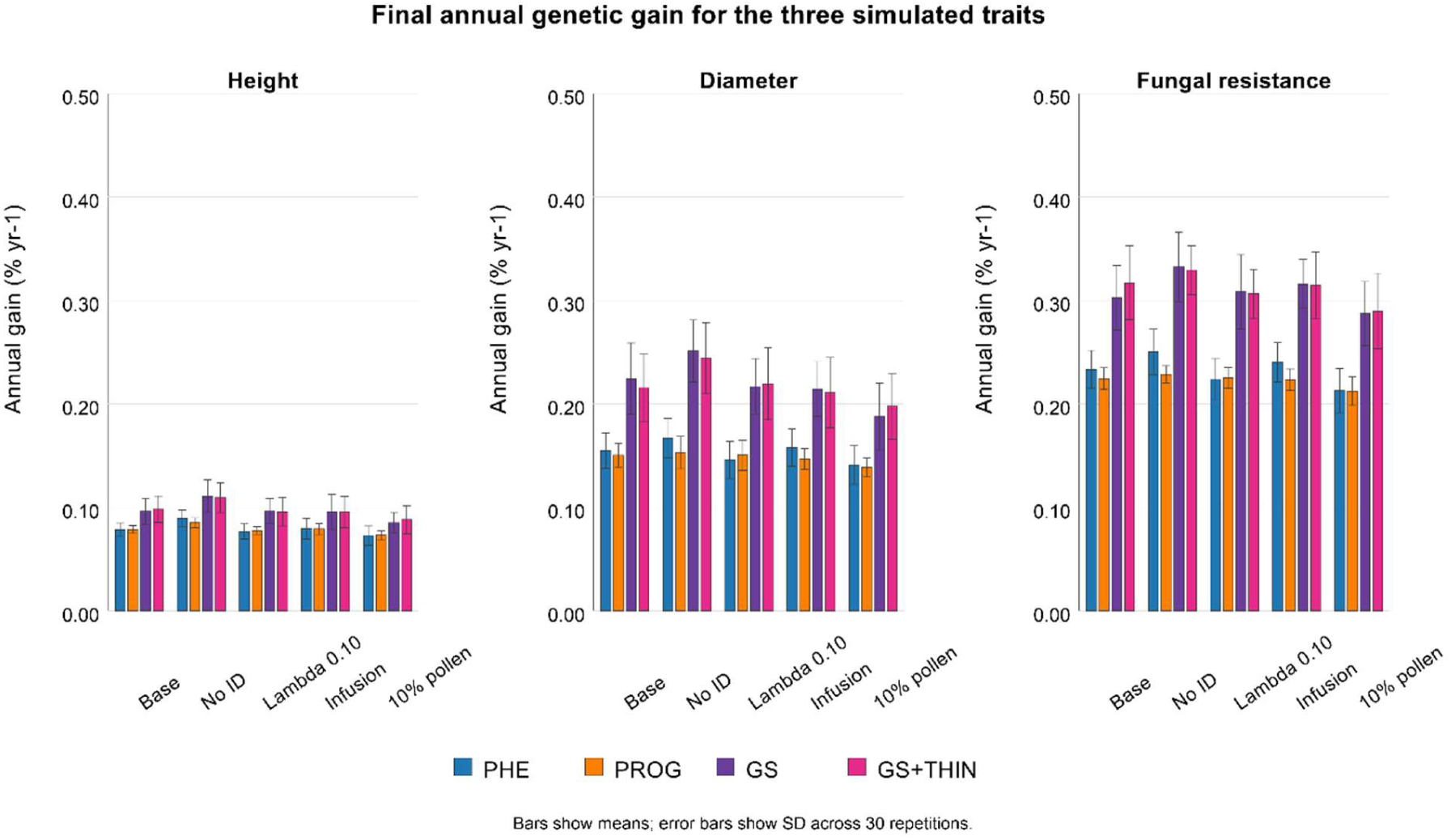
Final annual genetic gain for the three simulated traits. Annualisation used 30 years per phenotypic-selection cycle, 45 years per progeny-test cycle, and 31 years for the first GS or GS+THIN cycle followed by 16 years for each later cycle.

#### Inbreeding, kinship, and ancestor diversity

The genomic strategies accumulated more relatedness than the conventional strategies (figure 4). Starting from mean kinship of approximately 0.013, final BASE kinship was 0.0323 for PHE and 0.0349 for PROG, compared with 0.0527 for GS and 0.0534 for GS+THIN. Corresponding final ancestor-diversity values were 169.4, 163.8, 146.7, and 147.3, respectively. The NO_ID sensitivity variant increased final kinship to 0.0582 for GS and 0.0574 for GS+THIN and reduced ancestor diversity to 142.4 and 143.7. Thus, the increased annual response obtained when inbreeding depression was ignored was accompanied by additional, although moderate, concentration of parental ancestry.

**Figure 4.**
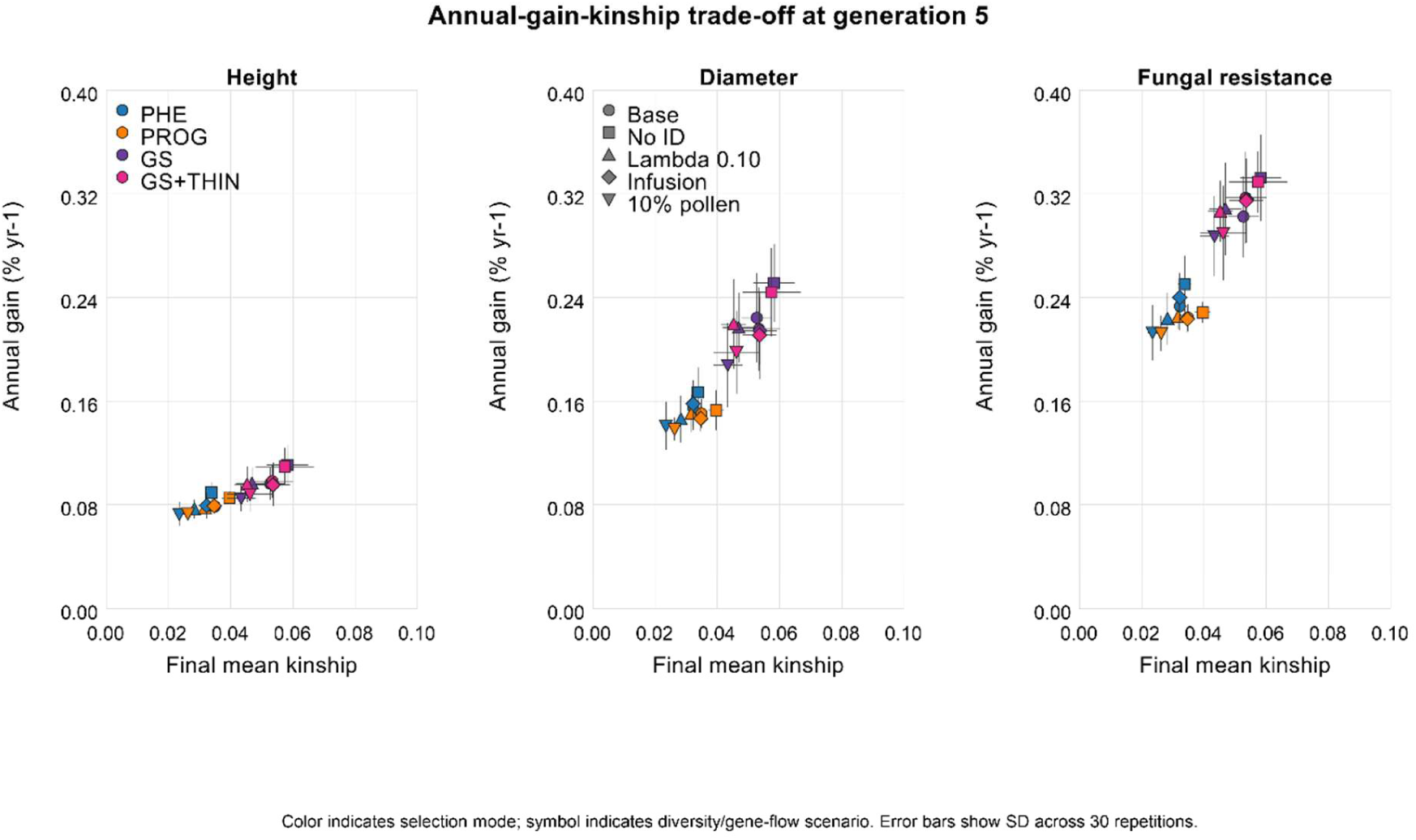
Annual-gain–kinship trade-off at generation 5 for the three simulated traits. Each point represents one selection-strategy × diversity-variant combination; favourable solutions combine high annual gain with low kinship.

The three active mitigation variants produced different trade-offs. For pure GS, external pollen reduced final kinship from 0.0527 under BASE to 0.0433. For GS+THIN, the λ = 0.10 kinship penalty reduced kinship from 0.0534 to 0.0452 and increased ancestor diversity from 147.3 to 156.2, while cumulative gains remained close to BASE at 9.09%, 20.85%, and 29.11% for height, diameter, and fungal resistance. Periodic infusion left GS+THIN kinship essentially unchanged at 0.0536 and increased ancestor diversity slightly to 149.2, with near-BASE gains. External pollen also lowered GS+THIN kinship to 0.0462, but reduced cumulative gains to 8.38%, 18.79%, and 27.51% and did not increase tracked ancestor diversity because new founder identities were not created. No mitigation option simultaneously maximised gain, prediction accuracy, and diversity. Complete diversity metrics and their graphical comparison are provided in Supplementary Table S9 and Figure S7.

#### Response dynamics and genomic-prediction accuracy

The BASE trajectories illustrated the different temporal profiles of the four strategies (Supplementary Figure S8). The first genomic cycle required 31 years, but subsequent cycles required only 16 years, so GS and GS+THIN completed five cycles within 95 years, compared with 150 years for PHE and 225 years for PROG. Both genomic strategies exceeded PHE and PROG in final BASE annual gain for all three traits. GS+THIN was highest for height and fungal resistance, whereas pure GS was highest for diameter. Differences between GS and GS+THIN were modest and trait dependent, so phenotypic thinning did not provide a uniform gain advantage.

Final genomic-prediction accuracy, defined as the correlation between PGBV and true breeding value, was 0.476 for GS and 0.490 for GS+THIN under BASE. Across all genomic strategy × diversity-variant combinations, final accuracy ranged only from 0.466 to 0.491, with the highest value for GS+THIN under POLLEN_10. These comparatively homogeneous accuracies show that realised differences among scenarios were driven more by cycle length, phenotypic thinning, inbreeding depression, and diversity management than by large differences in prediction accuracy. Accuracy alone did not determine the realised outcome. Complete accuracy statistics are provided in Supplementary Table S10 and Figure S9.

## Discussion

### Long-term progeny trials as genomic reference populations

The Drewin and Eiserbude trials demonstrate both the scientific value and the practical limitations of converting historical field tests into genomic reference populations. Their repeated measurements, mature age, and representation of controlled-cross, mixed-pollen, and seed-orchard material provide information that cannot be recreated rapidly. At the same time, two genomic quality-control layers were required before final inference: offspring-level likelihood-based parentage auditing and KING-based near-duplicate exclusion. The final population of 1,651 trees is therefore more biologically defensible than the unfiltered recorded pedigree.

This finding is consistent with operational pine studies in which genomic relationships improved the description of realised family structure, genetic parameters, and individual ranking (Tambarussi et al. 2025). It is also relevant to open-pollinated and mixed-mating material, for which simple half-sib assumptions can misrepresent non-random relationships. Empirical and simulation work in Eucalyptus has likewise shown that ignoring family structure and inbreeding can create gaps between predicted and realised gain (Araujo et al. 2025). For the German Scots pine breeding programme, the PiSy50k data therefore provide both a prediction resource and an enduring quality-control layer for crosses, seed lots, DNA samples, and future reference-population updates.

Individual-level parentage classification was less wasteful than excluding entire heterogeneous progenies. It retained supported offspring while removing records whose recorded pair was not the best-supported candidate, and it treated mixed-pollen families as mother-known, father-unknown material rather than failed controlled crosses. Retaining the KING duplicate screen additionally prevented repeated genotypes from being over-represented in variance estimation, cross-validation, and candidate ranking.

### Trait-and age-dependent value of genomic relationships

On identical parentage-green no-duplicate records, the genomic advantage was moderate, trait dependent, and age dependent. GBLUP increased heritability and both validation metrics for diameter and volume from age 20 onwards. At age 35, the leave-progeny-out gains were 0.033 for diameter, 0.016 for height, and 0.035 for volume. Height was more mixed: ABLUP produced higher heritability at several juvenile and mid-age assessments, whereas h35 heritability was essentially identical and GBLUP showed small positive prediction gains. This is broadly consistent with Calleja-Rodriguez et al. (2020), who found similar genomic and pedigree performance for many Scots pine traits and emphasised the value of earlier selection.

The results also agree with the wider forest-tree evidence that adding markers does not automatically produce large improvements. A meta-analysis across 28 genomic-selection studies found that predictive accuracy was more strongly associated with training-set size and breeding-population relatedness than with marker numbers once genome coverage was adequate (Beaulieu et al. 2024). The present marker panel is therefore unlikely to be the principal limitation. Greater gains should be sought by increasing the number of well-phenotyped and well-connected families, balancing family representation, improving phenotype quality, and maintaining pedigree links across generations rather than by increasing SNP density alone.

Heritability estimates from ABLUP and GBLUP should not be interpreted as competing measurements of a fixed biological constant. They depend on the relationship matrix, sample composition, fixed effects, trait scale, and connectedness represented in the data. The present matched comparison deliberately used identical individuals, phenotypes, folds, and fixed effects, so its differences isolate the effect of expected pedigree relationships versus realised genomic relationships. ABLUP nevertheless remains a strong operational baseline and can exploit larger phenotypic populations where complete genotyping is not available.

The ssGBLUP robustness analysis did not change this interpretation. Integrating A and G in H produced stable models and occasional small prediction gains, but ssGBLUP did not consistently outperform the better matched model and yielded lower age-35 heritabilities. Because all phenotype-bearing trees in this comparison were genotyped, the analysis tested relationship-matrix integration rather than the principal operational advantage of single-step evaluation— adding phenotypes from non-genotyped relatives. Its performance also depended on the chosen genomic blend and update weight. ssGBLUP is therefore best regarded here as a successful technical extension rather than a replacement for the main ABLUP–GBLUP comparison.

### Validation design and transfer across families and generations

Validation design was central to interpretation. Random-individual folds retained relatives of validation trees in training and measured interpolation within the represented population. Leave-progeny-out validation was more demanding because complete progenies were withheld. Positive genomic deltas for mature diameter, height, and volume under this conservative design strengthen the evidence that GBLUP contributes beyond simple interpolation among recorded sibs.

The unique contribution of genomic relationships is the estimation of Mendelian-sampling deviations among relatives. In maritime pine, Papin et al. (2024) showed that satisfactory overall prediction can coexist with weak within-family accuracy when families are represented by too few genotyped siblings. The present within-family analysis likewise produced weak correlations, especially for ABLUP, and GBLUP correlations remained close to zero for the age-35 traits. Most useful signal therefore arose from between-family differences and realised relationships across the population, and the data do not justify strong claims about precise full-sib ranking.

Recent independent and multi-generational studies show that cross-generation prediction is feasible, but they also clarify the conditions required. Hayatgheibi et al. (2025) obtained useful cross-generational predictions for Norway spruce wood properties with independent validation, while emphasising context-specific models and the need to account for age and environment. In loblolly pine, Isik et al. (2026) reported high genomic accuracies and substantially greater gain per year in a well-connected two-generation programme; prediction accuracy was strongly related to the mean relationship between training and validation populations. These studies support the direction of the German programme, but they also argue against treating the present 1990 trials as a static reference population. Each new cycle should contribute family-balanced genotypes and high-quality phenotypes so that the prediction model remains connected to the current selection candidates.

### Early selection, site effects, and expansion of the breeding objective

The age-age results identify age 20 as a plausible intermediate decision point for growth. Diameter and volume at age 20 were strongly correlated with age-35 performance, whereas the very early height relationships were unstable when site means were pooled. Earlier Scots pine research likewise found that short growth-chamber tests did not necessarily predict 20-year field performance, even when competition was manipulated to mimic later stand conditions (Jansson et al. 1998). Together with the time trends reported by Haapanen (2001), this supports an evidence-based rather than fixed definition of early selection age: the earliest operational age should be trait specific and should be validated within environments and breeding populations.

The moderate progeny-mean correlations between Drewin and Eiserbude showed shared ranking but incomplete equivalence of site performance. Site-specific heritabilities and prediction results varied in magnitude, particularly for diameter and volume. Because the full parentage-aware multi-environment variance partition was numerically unstable, it was not retained. The defensible conclusion is therefore limited to site sensitivity: combined breeding values should be interpreted together with trial-specific performance, and stronger G×E claims require additional trials or more stable covariance models.

Multi-trait and high-throughput phenotyping offer a practical route to improve both prediction and environmental interpretation. In lodgepole pine, multi-trait models increased predictive accuracy and reduced bias across growth, wood, pest, drought, and defence traits (Cappa et al. 2022). Multivariate genomic models also improved prediction of spring frost tolerance in Norway spruce by using genetically correlated bud-burst information (Aro et al. 2025). For Scots pine, LiDAR height assessment has already produced selections closely aligned with conventional measurements (Liziniewicz et al. 2025), while hyperspectral traits showed heritable variation and stability across sites and seasons (Provazník et al. 2025). These technologies could expand the German reference population at lower marginal phenotyping cost and provide adaptive traits that are difficult to measure repeatedly by conventional field methods.

The hypothetical fungal-resistance trait in the simulations should be interpreted in this broader context, not as a parameter estimate from Drewin and Eiserbude. Resistance to Scots pine blister rust has shown moderate narrow-sense heritability and high cross-site genetic correlation without an adverse genetic correlation with height or vitality (Persson et al. 2024). Drought-related root and canopy traits also displayed moderate to high heritability in Scots pine, although breeding populations showed reduced variation for some traits (Gil-Muñoz et al. 2025). These studies demonstrate that resilience traits can be integrated into Scots pine improvement, but the simulated heritability and major-gene architecture must be replaced by trait-specific empirical parameters before the model is used to recommend operational selection for disease or drought resistance.

### Breeding-cycle length and the role of phenotypic thinning

The simulations reinforce the central principle that tree-breeding strategies should be compared in gain per unit time rather than gain per generation alone (Grattapaglia and Resende 2011, Grattapaglia et al. 2018). Progeny testing produced the largest cumulative response over five generations, but the assumed 45-year cycle reduced its annualised advantage. Genomic strategies had to absorb a 31-year first cycle to establish the initial reference population, after which 16-year cycles became possible. Under the BASE assumptions, both genomic strategies achieved higher annual gains than phenotypic selection and progeny testing for all three traits: GS+THIN was highest for height and fungal resistance, whereas pure GS was highest for diameter. Their operational advantage therefore arose primarily from cycle shortening rather than superior cumulative gain per generation or a dramatic prediction-accuracy advantage. The strategy ranking depended on trait architecture, prediction accuracy, cycle length, and inbreeding depression; it was not an intrinsic property of the selection method.

Pure genomic selection and GS+THIN performed similarly, with trait-specific differences. Phenotypic thinning modestly improved height and fungal-resistance response, whereas pure GS was slightly higher for diameter. The operational rationale for the two-stage strategy is therefore not a uniform simulated gain advantage, but the retention of phenotypic verification before final reproductive selection. Genomic prediction reduced 2,000 seedlings to a manageable shortlist, after which field or orchard phenotypes could exclude candidates that failed minimum quality, health, or adaptation criteria. This is operationally credible for a long-lived species and preserves a mechanism for updating the reference population without assuming that early genomic predictions are error free.

Phenotypic thinning nevertheless imposed two successive selection filters, but its final BASE kinship was nearly identical to that of pure GS (0.0534 versus 0.0527). Both genomic strategies retained more relatedness than phenotypic selection and progeny testing. The use of thinning should therefore not be interpreted as support for simply increasing selection intensity or as an automatic diversity safeguard. The promising operational element is the division of labour between genomic and phenotypic information: genomic prediction determines which candidates merit further investment, while field and orchard phenotypes verify realised performance and maintain model relevance. Costed simulations should next vary the shortlist size, phenotyping intensity, and the age and precision of the thinning measurement.

### Diversity management determines long-term sustainability

The clearest diversity warning from the simulations was the consistent increase in relatedness under genomic selection, although the broadened burn-in population kept absolute values moderate. Under BASE, final mean kinship reached 0.0527 for GS and 0.0534 for GS+THIN, compared with 0.0323 for phenotypic selection and 0.0349 for progeny testing. The genomic strategies retained approximately 147 tracked ancestors, compared with 164–169 for the conventional strategies. Removing inbreeding depression produced the highest annual gains, but also the highest genomic-strategy kinship, approximately 0.058, and the lowest ancestor diversity, approximately 142–144. The NO_ID scenario is therefore useful only as an upper-bound sensitivity analysis. It illustrates how a breeding strategy can appear efficient when the biological and deployment consequences of relatedness are omitted.

Empirical studies in conifers demonstrate that these consequences can be substantial and may affect reproduction, survival, and growth at different life stages. In Scots pine, self-pollinated progenies had only 18.3% mature seed compared with 70.9% in open-pollinated progenies, while inbreeding depression for post-germination survival up to age 23 ranged from 0.62 to 0.75; cumulative inbreeding depression across the life cycle reached 0.90–0.94 (Koelewijn et al. 1999). In maritime pine, an inbreeding coefficient of F=0.75 was associated with reductions of 27% in height, 37% in stem circumference, 63% in bole volume, and 89% in female fertility, with depression in annual height growth becoming stronger during unfavourable years (Durel et al. 1996). In loblolly pine, increasing relatedness reduced height by up to 21% and stem volume by up to 33% at age nine (Ford et al. 2015), while in radiata pine increasing inbreeding reduced diameter and survival, with depressions of 19% and 11%, respectively, at F=0.75 (Wu et al. 1998). These findings indicate that increased kinship can diminish realised genetic gain through reduced seed production, survival, growth, and reproductive output, even when predicted breeding values remain high. Diversity management should therefore be treated as an integral component of recurrent genomic selection rather than as a secondary constraint.

The mitigation variants exposed different mechanisms rather than a single solution. External pollen reduced kinship particularly for pure GS, whereas the λ = 0.10 penalty produced the lowest GS+THIN kinship and retained the most tracked founder ancestry among the genomic mitigation variants. Uncontrolled background pollen remains difficult to translate into a predictable breeding contribution and, in the current implementation, did not create new tracked founder identities. Empirical studies show that background pollen contamination in pine seed orchards can vary by more than an order of magnitude, depending particularly on orchard age, isolation, and the strength of the orchard’s own pollen cloud. In a mature Swedish Scots pine seed orchard, contamination was only 4.89–7.29% across three consecutive pollination seasons, whereas an advanced Scots pine orchard showed 52% external paternity (Torimaru et al. 2009, Funda et al. 2015). A broader Swedish study found that background pollen contamination in Scots pine seed crops declined from 87% in a young orchard to 12% in a mature orchard (Heuchel et al. 2022). Similarly, 85.7% of viable seed in an 11-year-old Turkish red pine (*Pinus brutia*) orchard was attributed to pollen from surrounding stands, leaving only 9% of the offspring derived from crosses among orchard clones; the expected genetic gain was estimated to be less than 57% of that predicted in the absence of contamination (Kaya et al. 2006). Thus, the 10% external-pollen level used in the present simulations is realistic for a mature, well-functioning orchard but lies near the lower end of published observations. In practice, background pollen may increase diversity and reduce relatedness, but it can also dilute genetic gain and introduce paternal contributions of uncertain breeding value or adaptation. Because external pollen did not create new founder identities in the present model, the simulations captured its kinship-reducing effect but not its full potential contribution to founder diversity.

Periodic infusion introduced known, unrelated material and improved some prediction and response outcomes, but the number and timing of immigrants were insufficient to reduce kinship consistently across selection strategies. Operational examples suggest that effective infusion is usually based on larger, pre-evaluated sets of germplasm rather than a small number of untested immigrants. In radiata pine, an Australian programme selected 210 elite trees representing all five native provenances for controlled crossing and possible infusion into existing breeding populations, with the aim of broadening the genetic base and introducing favourable alleles for growth, wood properties, climatic adaptation, and pest or disease resistance (Gapare et al. 2012). In the Argentine loblolly pine breeding programme, a later test series incorporated two unrelated infusion sets—15 first-generation selections and 83 second-generation half-sib families from several geographic and breeding sources—within a series of 180 half-sib families intended to enhance genetic diversity (Belaber et al. 2025). In fourth-cycle Chinese fir, external superior trees were incorporated through genetic infusion, and the resulting 233-parent breeding population retained observed and expected heterozygosities of 0.215 and 0.233, respectively, with weak differentiation among breeding generations and germplasm origins (Jing et al. 2023). A subsequent restructuring of this population established a 50-member core population with an inbreeding coefficient close to zero and a 183-member main population with an effective population size of 108 (Zhao et al. 2023). These examples support infusion as an operational method for broadening advanced breeding populations, but they do not isolate its causal effect from selection history, mating design, and population restructuring. Accordingly, the ten immigrants introduced every second generation in the present simulations should be interpreted as a modest infusion treatment. In the revised simulations, the fixed λ = 0.10 kinship penalty retained more founder ancestry with little change in cumulative gain; it was therefore not overly restrictive and provided a favourable trade-off among the tested GS+THIN variants. A dynamically adjusted penalty or an optimal-contribution approach that balances breeding value against coancestry may nevertheless provide a more robust compromise between short-term gain and long-term diversity.

A more defensible next step is optimum contribution selection combined with planned mate allocation rather than a single fixed penalty. In a Scots pine application, optimum contribution selection achieved 8-30% more genetic gain than standard restricted selection at the same coancestry constraint (Hallander and Waldmann 2009). Such methods can assign unequal but controlled parental contributions and directly trace the gain-diversity frontier. They are especially compatible with genomic relationships, which quantify Mendelian sampling and realised coancestry among candidates more precisely than pedigree expectations. Future SNPscan_breeder scenarios should therefore compare optimum contributions, family contribution limits, minimum-coancestry mating, and controlled infusion using the same annual-gain accounting.

### Implications for the German Scots pine breeding programme

The empirical and simulation results favour an integrated transition rather than replacement of conventional breeding. The parentage-audited Drewin and Eiserbude trials can provide an initial genomic reference population. An operational pilot should genotype a large, family-balanced seedling cohort, predict growth and quality, and use genomic values to define a shortlist for field or seedling-orchard assessment. Final parent selection should combine genomic breeding values, observed performance, site response, minimum trait standards, and an explicit coancestry constraint.

Reference-population updating should be treated as a permanent programme function. A connected subset of each generation should be phenotyped across representative environments, with deliberate representation of all parental groups rather than only the highest-ranked families. Once marker density is sufficient, the number, quality, and connectedness of phenotyped individuals are likely to contribute more to prediction than denser genotyping (Beaulieu et al. 2024). The field network should therefore be redesigned to serve both selection and model maintenance, including rapid phenotyping of juvenile growth and adaptive traits, mature validation, and monitoring of genotype-by-environment interaction.

The simulation results also provide a decision criterion for programme governance. A genomic strategy should be adopted only if its realised annual gain, cost, and diversity trajectory improve on the conventional benchmark. This requires reporting not only prediction correlation and mean gain, but also family representation, mean kinship, effective population size, ancestor diversity, and deployment diversity. Periodic recalibration against realised field performance will be essential because climate, health threats, breeding objectives, and the genetic composition of the candidate population will change over the decades represented by the simulations.

### Limitations and priorities for further work

The empirical analyses were constrained by the available family structure, the non-random set of genotyped survivors, and the reconstruction of historical parents as latent rather than directly genotyped individuals. The 500-SNP audit provides strong comparative evidence but direct parental genotypes would be preferable. Stem form was analysed as a numeric score rather than with a threshold model. The full G×E variance partition was unstable, and the within-family results did not support precise Mendelian-sampling prediction. These limitations support staged implementation and continued validation rather than immediate replacement of pedigree evaluation. The ssGBLUP robustness analysis used a fixed 10% A22 blend and λ = 0.5 and included no additional non-genotyped phenotype records, so alternative tuning and a genuinely mixed genotyped–non-genotyped population remain to be evaluated.

The simulations represent a single burn-in population and 12-chromosome genome realisation, one randomly sampled 10,000-SNP non-causal prediction panel, one population size, trait architecture, selection intensity, and time-cost structure. Absolute kinship and ancestor-diversity values are sensitive to founder composition and the burn-in process. Height and diameter were informed by the empirical trial scales, whereas fungal resistance was hypothetical. The widened upper trait thresholds avoided artificial ceilings but remain configured plausibility bounds rather than biological maxima. Inbreeding depression was represented as a linear trait-level penalty rather than through dominance, deleterious alleles, or explicit genetic load. The simulations covered five recurrent cycles, used one kinship-penalty strength, and did not attach economic costs or uncertainty distributions to the breeding-cycle assumptions. The resulting rankings are scenario comparisons, not forecasts of realised gain.

Priorities are independent validation in the next breeding generation, enlargement and family balancing of the reference population, stable multi-environment and multi-trait modelling, direct genotyping of available parents or archival material, and a costed GS+THIN pilot. Simulation sensitivity should vary reference-population size, number of selected parents, shortlist size, phenotype age and cost, prediction decay across generations, optimum-contribution constraints, mating design, and the frequency and source of infusion. Selection-index weights, genetic-correlation files, burn-in populations, genome maps, software versions, and random seeds should remain archived for reproducible updating.

## Conclusions

Dense SNP data, likelihood-based parentage auditing, and duplicate filtering converted the Drewin and Eiserbude trials into a more informative and quality-controlled breeding resource. Across the longitudinal measurements, GBLUP improved prediction most consistently for diameter and volume from age 20 onwards. Height showed a more mixed pattern, with nearly identical ABLUP and GBLUP heritability at age 35 and smaller positive genomic prediction gains. Strict sensitivity analyses confirmed the mature-trait conclusions, while site-specific and within-family results showed that genomic advantages remain moderate and context dependent. The trials can form the first reference population for German Scots pine genomic selection if future training data are collected with balanced family and site representation. ssGBLUP fitted stably but did not consistently outperform the simpler matched models.

The simulations showed that the main opportunity from genomic selection is faster recurrent selection rather than superior gain per generation. Under BASE, both genomic strategies delivered higher annual gains than the conventional strategies for all three traits, while pure GS and GS+THIN differed only modestly and in a trait-dependent manner. Phenotypic thinning remains operationally credible because it provides field verification and a model-updating step, not because it uniformly increased simulated gain. The genomic strategies still produced higher kinship than phenotypic selection and progeny testing, but the λ = 0.10 penalty improved founder retention with little loss of response. Operational implementation should therefore retain field testing, update prediction models in every generation, and use optimum parental contributions and planned mating to constrain coancestry and preserve founder diversity. A staged genomic-shortlisting and GS+THIN pilot with explicit cost, annual-gain, prediction, and diversity targets is the most defensible next step; complete replacement of progeny testing is not supported by the present evidence.

## Acknowledgments

We thank Ute Straßburg-Käßler and Stefan Jencsik for technical assistance with field measurements, sampling, DNA extraction, and DNA quality control. Further we are thankful to our gardeners and the staff of the respective forest districts for long term maintenance of the trials This research was funded by the German Federal Ministry of Agriculture, Food and Regional Identity (BMLEH) via the funding agency FNR, grant number 2223NR023X. We used AI (ChatGPT version 5.5) to check and optimise r-scripts for the data analysis and to polish the language.

## Data Availability Statement

In case of acceptance the SNP-data and the phenotype-data of the two trails as well as the input data and parameter settings of the simulations will we assessable at Zenodo. The current version of the simulation model SNPscan_breeder 3.1 and a user’s manual will be available at https://www.thuenen.de/en/institutes/forest-genetics/software.

